# Neuronal correlates of sleep in honey bees

**DOI:** 10.1101/2024.10.11.617548

**Authors:** Sebastian Moguilner, Ettore Tiraboschi, Giacomo Fantoni, Heather Strelevitz, Hamid Soleimani, Luca Del Torre, Uri Hasson, Albrecht Haase

## Abstract

Honey bees *Apis mellifera* follow the day-night cycle for their foraging activity, entering rest periods during darkness. Despite considerable research on sleep behaviour in bees, its underlying neurophysiological mechanisms are not well understood, partly due to the lack of brain imaging data that allow for analysis from a network- or system-level perspective.

This study aims to fill this gap by investigating whether neuronal activity during rest periods exhibits stereotypic patterns comparable to sleep signatures observed in vertebrates. Using two-photon calcium imaging of the antennal lobes (AL) in head-fixed bees, we analysed brain dynamics across motion and rest epochs during the nocturnal period. The recorded activity was computationally characterised, and machine learning was applied to determine whether a classifier could distinguish the two states after motion correction. Out-of-sample classification accuracy reached 93%, and a feature importance analysis suggested network features to be decisive. Accordingly, the glomerular connectivity was found to be significantly increased in the rest-state patterns. A full simulation of the AL using a leaky spiking neural network revealed that such a transition in network connectivity could be achieved by weakly correlated input noise and a reduction of synaptic conductance of the inhibitive local neurons (LNs) which couple the AL network nodes. The difference in the AL response maps between awake- and sleep-like states generated by the simulation showed a decreased specificity of the odour code in the sleep state, suggesting reduced information processing during sleep. Since LNs in the bee brain are GABAergic, this suggests that the GABAergic system plays a central role in sleep regulation in bees as in many higher species including humans. Our findings provide the first evidence that sleep-related network modulation mechanisms may be conserved throughout evolution, highlighting the bee’s potential as an invertebrate model for studying sleep at the level of single neurons.

## 1. Introduction

The sleep phenomenon in invertebrates is not yet well understood (Siegel, 2008). A promising model for its investigation is the honey bee *Apis mellifera*. An early study of bee behaviour during an entire life cycle showed that bees spend a large part of their time in what was termed rest phases (Lindauer, 1952). Bees are commonly referred to as sleeping when they assume a typical body position with characteristically hanging antennae (Kaiser, 1988). During these phases, responsiveness to external stimuli is reduced (Eban-Rothschild & Bloch, 2008). While young workers, dedicated to in-hive activity, show an irregular pattern of sleep phases, foragers’ sleep is strongly linked to the circadian rhythm (Yerushalmi et al., 2006). Sleep in bees is found to improve memory consolidation (Hussaini et al., 2009; Zwaka et al., 2015), while sleep deprivation affects memory extinction (Beyaert et al., 2012), reduces waggle dance precision (Klein et al., 2010), and is later compensated by increased sleep (Stefan Sauer et al., 2004). Furthermore, honey bee sleep can be influenced by neuroactive pesticides, such as neonicotinoids (Tackenberg et al., 2020; Tasman et al., 2020) and herbicides like glyphosate (Vázquez et al., 2020). However, little is known about the dynamic neural transition mechanisms underlying sleep.

So far, the only neurophysiological signature of sleep reported in the bee brain is a circadian modulation of electrophysiological responses of single neurons in the optic lobes of foragers (Kaiser & Steiner-Kaiser, 1983). However, an expression of the circadian clock gene PER was found throughout the bee brain (Bloch et al., 2003). This suggests that sleep impacts whole-brain activity in a measurable, but yet non-understood way. This includes the primary olfactory processing centers, the antennal lobes (ALs), which have so far been ignored by sleep studies in insects (Helfrich-Förster, 2018). In the ALs, olfactory receptor neurons (ORNs) from the antennae converge into 160 nodes called glomeruli, each receiving input from a single type of odour receptor. These glomeruli are interconnected via inhibitory local neurons (LNs) and project into higher brain centers via projection neurons (PNs). Odour information is coded in spatio-temporal activity patterns in the PNs (Paoli et al., 2016, 2018). To gain a better understanding of the neuronal basis of sleep in bees, and given that the bee olfactory system has been shown to be functional during sleep (Zwaka et al., 2015), we chose to record neuronal activity patterns in the ALs, both at the network level and at the level of univariate time series.

In this study, we aimed to provide the first study of the neural correlates of sleep in the bee brain and explore its functional implications. By combining functional brain imaging, machine learning, and computational modelling, we applied an integrated approach to study sleep at both experimental and theoretical levels. We used extended recordings of two-photon calcium imaging to monitor the spontaneous activity in the PNs during phases of distinct motor activity to quantify functional changes associated with sleep.

To this end, we implemented a hybrid data-driven approach via machine learning and a hypothesis-driven approach. Specifically, we raised three hypotheses: (1) Rest phases will be associated with reduced information processing at the network level, significantly different from the active state correlates, as observed also in the human brain (Alkire et al., 2008; Tononi & Massimini, 2008). (2) Reduced information processing in the antennal lobe should manifest in a reduction of contrast and sparsening of glomerular activity at the output. (3) The reduction of function in the antennal lobe could be generated by a change in glomerular coupling via the LNs. By testing these hypotheses, we aim to gain new insights into the neural correlates of sleep in invertebrates and to understand to what extent these brains with only one million neurons allow the future study of sleep-related effects.

## 2. Methods

### 2.1. Experimental procedures

#### 2.1.1. Insect preparation

Forager honey bees (*Apis mellifera*) from outdoor hives were captured in the morning (between 9 a.m. and 11 a.m.) at a feeder using small plastic containers. The preparation then closely followed a standard procedure (Paoli et al., 2017). Briefly, after immobilization by cooling the temperature of the containers, bees were transferred into a Plexiglass mount (**Fig. 1a**), where the head was fixed with soft dental wax (Deiberit 502, Siladent). A small window was cut into the head cuticula, and glands and tracheae were gently moved aside to clean the dye injection site. A borosilicate glass needle was used to inject the calcium-sensitive dye Fura-2-dextran (ThermoFisher Scientific). The injection site (always in the left brain side to avoid lateralization effects (Haase, Rigosi, Frasnelli, et al., 2011) was at the intersection of the lateral and medial antenno-protocerebral tracts between medial and lateral calyxes of the mushroom body (Paoli et al., 2016). The cuticula was then closed; the bees were fed to satiety with 50% sucrose/water solution and kept in a dark, moist place for at least 6 h to allow for dye uptake. In the afternoon, the bees were prepared for microscopy by re-opening the cuticula, removing glands and tracheae from the imaging site (the left AL), and covering the brain with a transparent two-component silicon (Kwik-Sil, WPI). A small (3 - 4 cm) transparent plastic sheet was placed on top of the bee’s head in order to isolate the dry antennae from the imaging window, where a large drop of water was added to immerse the microscope objective. To compensate for the evaporation of the immersion liquid during the night, an automated refill system was constructed, which detected the absence of water via the current drop between two electrodes. This triggered an automated syringe to refill the reservoir.

**Figure 1.**
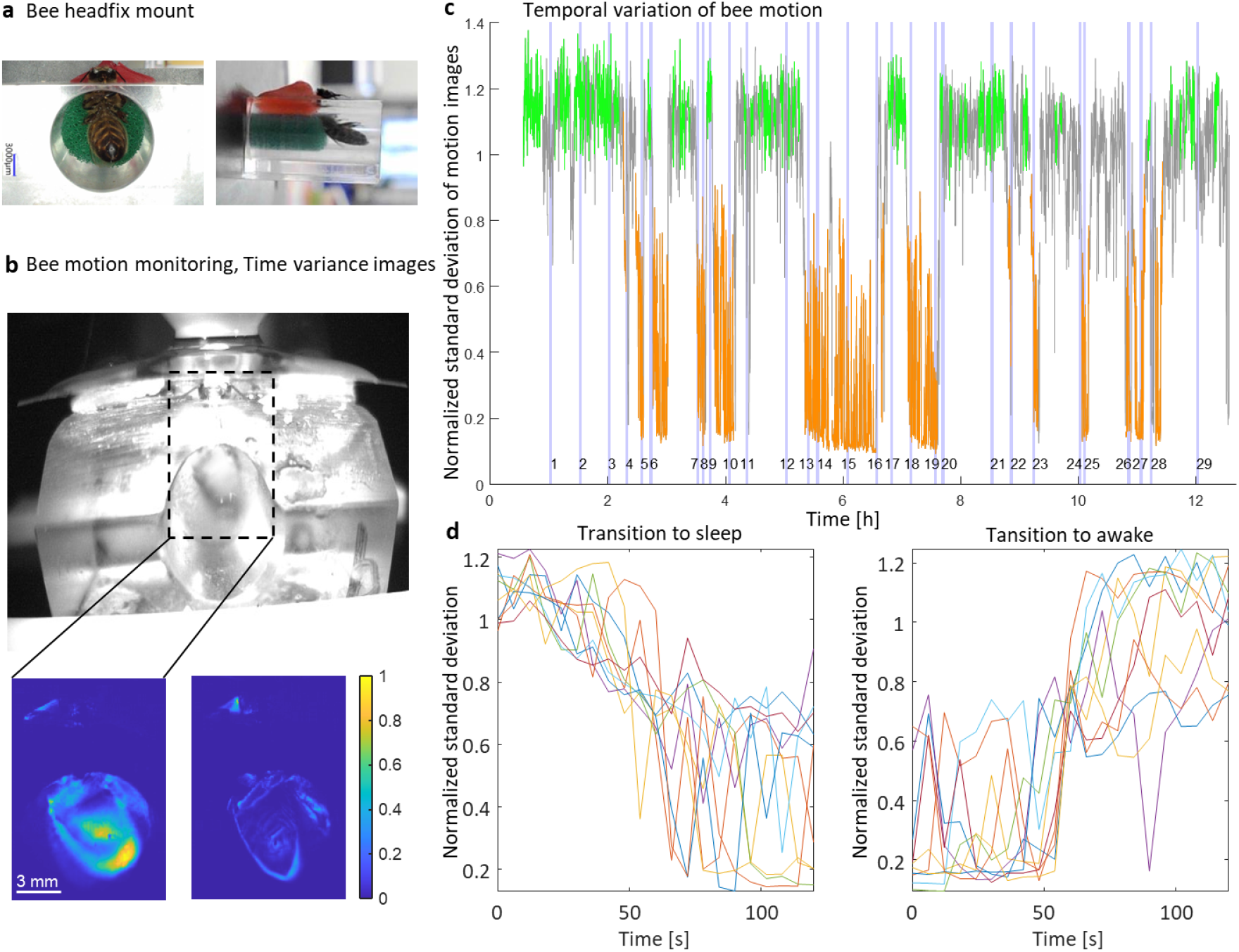
Trace of bee motion. **(a)** Honey bee in the plexiglass mounting block under two perspectives, with red dental wax to fix the head and a green sponge to gently fix the upper body. **(b)** above: Frontal camera view of the mounted bee under the microscope; below: Standard deviation of the intensity of each camera pixel during an interval of 200 frames, revealing a period of frequent trunk movement (left) and a resting phase (right). **(c)** Running standard deviation (200 frames) averaged over the entire camera image and normalised to an initial value. This measure of body motion alternates between states of strong motion (green) and resting states (orange), with each state typically extending from multiple minutes to over one hour. Intervals that could not clearly be classified as one of the two states are shown in grey. Light blue vertical bars mark the two-photon imaging periods of 2.5 min. In addition to regular imaging sessions every 30 min, recordings were triggered by the detection of a change in the state of motion with respect to the previous recording session. The motion images in (b) correspond to imaging session 2 (awake) and session 5 (rest). **(b)** Detailed analysis of the transition phases of the extended sleep periods in (c).

#### 2.1.2. Body motion video recording

The bees’ motion during imaging was recorded via an infrared camera (Blackfly BFLY-PGE-12A2M, FLIR), placed 15 cm in front of the bee in the microscopy mount, at a frame rate of 10 Hz. An illumination invisible to both the bee eye and the microscope detectors was provided by an infrared LED (*λ* = 850 nm). Bees were habituated to the setup for at least 1h, then a video of 30 minutes was recorded as a reference for the active-state motion (calculated from the standard deviation (*STD*_ref_) of each camera pixel’s time series over 200 frames, averaged over area and recording time). In agreement with previous circadian rhythm studies (Kaiser & Steiner-Kaiser, 1983), bees were never found resting before 9 p.m.

#### 2.1.3. Calcium imaging

The two-photon imaging platform (Ultima IV, Bruker) consisted of a Ti:Sa laser (Mai Tai Deep See HP, Spectra-Physics) tuned to 780 nm for fura-2 excitation, illuminating an epi-fluorescence microscope with a water immersion objective (10×, NA 0.3, Olympus). Fluorescence was recorded via a photomultiplier (Hamamatsu) through a 525 ± 35 nm filter (Chroma). A field of view of 280 × 280 µm^2^, resolved in 128 × 128 pixels, allowed to simultaneously image between 12-25 glomeruli. The image acquisition rate was approximately 10 Hz in regular scanning. A laser power of about 4 mW (after the objective) showed an optimal balance between signal-to-noise ratio and photo-damage, the latter manifested in the bee lifetime. For one bee, scanning via a resonant piezo mirror allowed acquisition at 120 Hz which produced a reduced signal-to-noise ratio that was partially compensated by increasing the laser power up to 14 mW. These powers are well below thresholds at which significant heating effects can be detected in brain tissue (~1.8°C /100 mW) (Podgorski & Ranganathan, 2016), and identical imaging parameters during sleep and wakefulness should rule out any differential effect of the laser. For all animals, two-photon recording sessions were triggered by real-time analysis of the bee’s motion state from the IR camera data and lasted 150 s each.

#### 2.1.4. Motion-triggered image acquisition

A MATLAB (MathWorks) script analysed the IR camera data every 150 s. Within these intervals, the pixel standard deviation (STD) was recomputed over every 200 frames collapsing over the entire field of view. Then, the temporal mean *STD*_mean_, the maximum *STD*_max,_ and the minimum *STD*_min_ were extracted and the ratios to the reference *STD*_ref_ were used as behavioural indicators to distinguish between active and resting states (details see below). The microscope was by default triggered every 30 min for 150 s imaging sessions. Additionally, to equilibrate the amount of data from the different motion states, motion classification was performed in real-time and whenever a shift of state with respect to the previous recording was detected, an additional imaging session was initialised. The light blue vertical lines in **Fig. 1c** show an example of the timeline of these imaging sessions for one animal.

#### 2.1.5. Post-hoc motion classification from video recordings

During the experiments, frontal videos were saved to allow a refined motion classification during post-processing and to section the calcium imaging data to ‘active’ and ‘resting’ conditions according to the motion states. The classification was based on the relative change of the image intensity standard deviation described above; the thresholds were optimised based on the fluctuations during the recording to robustly distinguish the two well-separated motion states (**Fig. 1c**). These thresholds were animal-dependent, not only due to behavioural differences between bees but also because of differences in the visible area of the bee body and the imaging angle. An example of classification thresholds for the recording visualized in **Fig. 1c** is for the resting state: 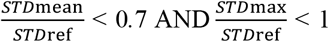 and for the active state: 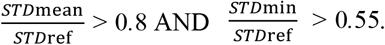 During a 150 s imaging session, bees could change state or fluctuations could compromise a clear classification; those data were excluded from further analyses.

To guarantee high data quality for the quantitative statistical analysis, experiments were only considered if they fulfilled each of the following criteria: a) images must allow for distinguishing individual glomeruli; b) bees must have survived the whole night; c) the real-time classification must have triggered a number of imaging sessions in both motion states that allow for a reasonable statistical analysis; d) the rest and active states must have been different enough to allow for a reliable classification according to the above criteria. After careful examination of all these criteria, we arrived at *n* = 9 subjects to be included in this study.

### 2.2. Image post-processing

#### 2.2.1. Brain motion correction

It is essential to apply motion correction in the absence of rigorous frame shift control since, during the analysis, motion might be mistakenly considered as a calcium-induced change in fluorescence. A Hidden Markov Model (HMM) (Dombeck et al., 2007) was used to correct for within-frame artefacts due to residual transverse brain motion. The HMM uses a maximum likelihood method to determine the most probable (*x,y*)-displacement between consecutive frames in a time series. The displacement was modelled as a first-order auto-regressive process in two dimensions, and the pixel values were assumed to result from linear scaling of photon counts that obey Poisson statistics (Kaifosh et al., 2013). To inspect the HMM motion correction performance, a calculation of the phase correlation between pixels of consecutive frames identified the relative shift before and after motion correction (**Fig. A.1**).

#### 2.2.2. Activity signal extraction

The fluorescence images (**Fig. 2a, A.4a**) were spatially smoothened using a 5 × 5 Gaussian kernel (**Fig. A.4b**). Along the time axis, data were partitioned into segments of 100 time points (ca. 10 s for normal scanning or 1 s for resonant scanning). This produced for each recorded bee approximately 50 - 60 segments in each activity state. Images were then normalised with respect to the average fluorescence during a time segment −Δ*F*/*F* (**Fig. A.4c**). Finally, they were thresholded between 0.5 and −0.5 (**Fig. A.4d**), which are the maximum calcium-induced fluorescence changes observed in previous experiments (Haase, Rigosi, Frasnelli, et al., 2011). The segmentation of individual glomeruli was guided by a regional homogeneity analysis (ReHo) that quantifies the local spatial autocorrelation for each pixel with its neighbours (Zang et al., 2004) (**Fig. 2b**). These ReHo maps show high values within glomeruli which decay at the border, allowing for easy identification and segmentation of glomeruli and for comparison of the coherence within individual glomeruli between active and rest phases. After segmentation of all identifiable glomeruli, the normalised activity was averaged over each glomerular ROI. This provides for each glomerulus a time series of the calcium-induced fluorescence change, a property that was shown to resemble the firing rate of the PNs in a glomerulus (Moreaux & Laurent, 2007).

**Figure 2.**
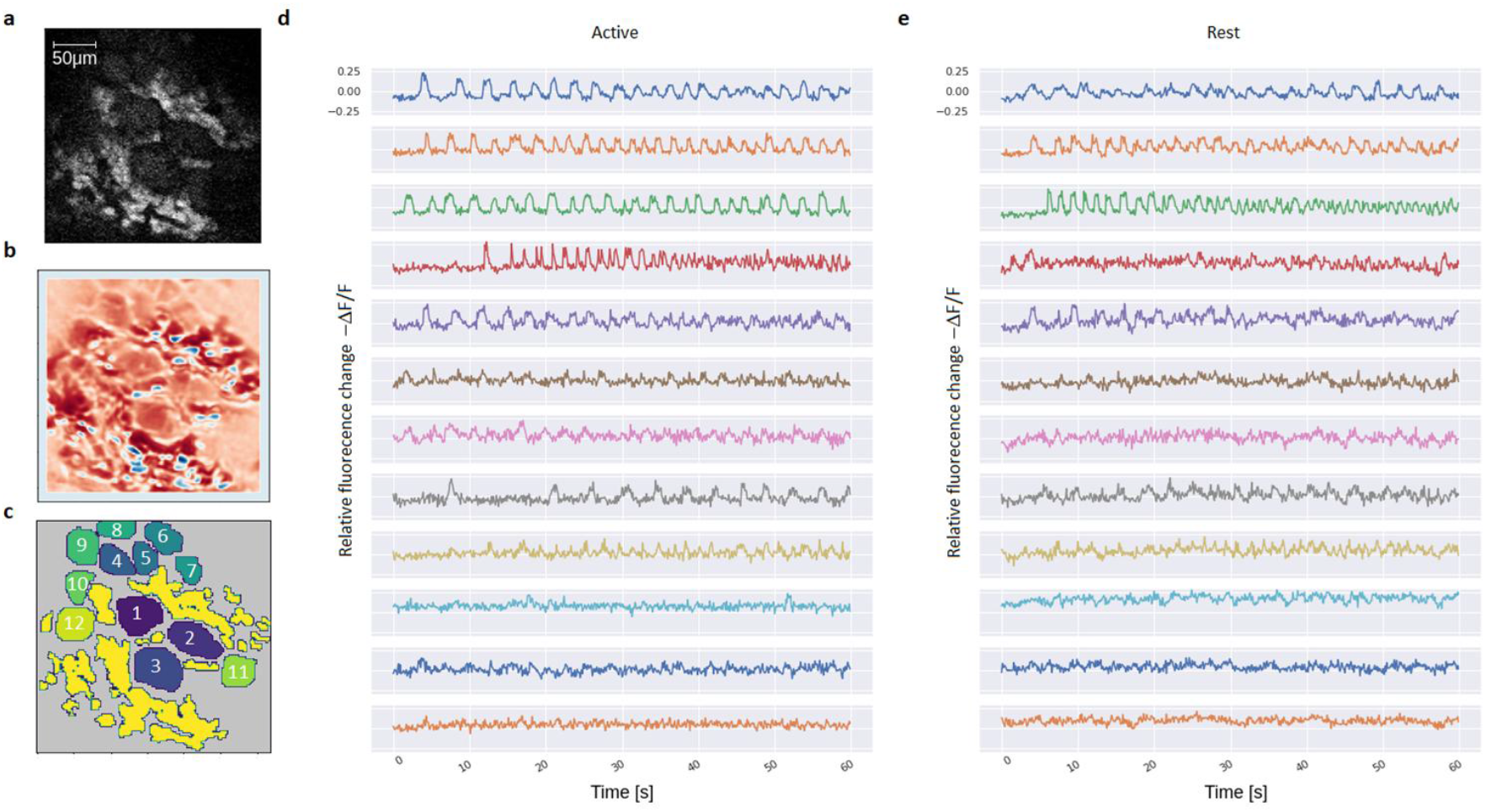
Glomerular signal extraction. **(a)** Fluorescence signal from a single focal plane through the antennal lobe, showing glomeruli (large, grey structures) and somata (small, white structures). **(b)** Regional Homogeneity (ReHo) analysis computes the similarity of the time series of each pixel and adjacent ones. The colour map changes with increasing homogeneity from blue to white to red. Large coherence is found at the centres of the glomeruli and decays towards their boundaries into white. The somata are located in the periphery and show no strong, synchronised activity, hence they appear in blue. **(c)** Individual glomerulus ROI map after ReHo-supported image segmentation. **(d)** Sample time series of 1 s of the normalised fluorescence variation (−Δ*F*/*F*) for an active motion state in each glomerulus in (c). **(e)** Sample time series of the normalised fluorescence variation for a rest state in each glomerulus in (c).

#### 2.2.3. Correlation analysis

To investigate the coupling between individual glomeruli, a cross-correlation analysis was performed between the activity time series for each pair of glomeruli within each bee (**Fig. 4a,b**). To exclude spurious correlations due to arbitrary fluctuations, surrogate data sets were produced by randomly shuffling segments of each glomerular time series. *T*-tests were performed to assess the difference between the glomerular pair correlation and the spurious correlation between the pair’s surrogate time series. Glomerular pairs with non-significant correlations were excluded from the calculation of the overall correlation in the glomerular network of each bee (**Fig. 4c**).

### 2.3. Machine learning methods

#### 2.3.1. Definition and analysis of features

In the spatial domain, not all of the 120 × 120 pixels per image were used for classification, but only those that were *a priori* known to provide the most relevant information. For this reason, feature quantification focused on signals originating from the glomeruli as obtained by image segmentation methods like the REHO analysis described above. For the classification, features characterising each time series segment were defined under three different categories: the general distribution of activity, the signal complexity in the time domain, and network features (for details see **Eq. (A.1-A.10**). The same processing pipeline was applied for each bee. First, time series were partitioned into segments of 100 time points length for each glomerulus in a bee (*i*.*e*. averaging across the pixels contained in a glomerulus). Then, to obtain the distribution moments and complexity features, each measure was first calculated for each glomerulus and then averaged to obtain a mean across glomeruli. For the connectivity features, graph theory features were computed from the functional connectivity matrix obtained from 100 time-intervals for each glomerulus. A statistical analysis of the features was performed, comparing them between active and rest states in individual bees and binning all bees. The Wilcoxon non-parametric rank test was used and the probabilities were corrected for False discovery rate (FDR).

#### 2.3.2. Training and Prediction

The features were Min-Max normalised but not scaled. The model was a Random Forest Classifier (RFC) (Pedregosa et al., 2011) that was trained to classify active from rest states. A standard RFC implementation on default hyper-parameters was used (*i*.*e*. 100 decision-tree estimators, 2 minimum sample splits per decision-tree branch, 1 minimum sample leaf for each branch, and 0 minimum weight fraction leaf). 80% of the segments were used as training data and 20% as unseen (out-of-sample) test data. Once the model was trained, predictions on the unseen data were produced and evaluated against the ground truth from the body motion data to produce a measure of accuracy.

#### 2.3.3. Interpreting the active-rest classification model

To obtain which features were the most relevant in each classification scheme, we performed a feature importance analysis. This process is based on the evaluation of the relative weights of the decision trees. Random forests consist of an ensemble of decision trees, in which every node in the decision tree is a condition on a single feature, designed to split the data into two sets so that similar response values end up in the same set. The Gini impurity (Menze et al., 2009) was employed to compute how much each feature decreases the weighted impurity in a tree. It is a measure of the likelihood of incorrect classification of an element if it were randomly classified according to the distribution of class labels in the data set. This allowed for measuring how each feature decreases the impurity of the split (the feature with the highest decrease is selected for the internal node). Finally, a measure of the feature importance was obtained from the average decrease of the impurity by each feature.

#### 2.3.4. Control by brain motion classification

To control whether motion artefacts contributed to the rest-active classification, a similar Random forest algorithm was trained to classify data segments into high and low brain motion states. This was done by using the sum over all *x*- and *y*-displacements during each segment after motion correction (**Fig. A.1b**). Rest and active states were pooled, and a median split into high and low brain motion segments was performed. A classifier was trained and tested identically to the one reported above for active vs. rest states.

### 2.4. Simulations

In a previous work, we demonstrated that a recurrent spiking neural network model of the antennal lobe topology could successfully reproduce odour response functions under the assumption of uniform synaptic coupling (Scarano et al., 2023). We used the powerful GPU-based neurocomputing tool GeNN (Knight et al., 2021) to simulate the antennal lobe network in the insect brain in their original complexity. Given the experimentally observed changes in glomerular coupling between sleep and wake states, we aimed to determine the simplest network modification that could account for these differences. The recurrent spiking neural network was modelled to closely resemble the biological system: neurons were organized in 160 glomeruli, each with 60 olfactory receptor neurons (ORN), 25 local neurons (LN), and 5 projection neurons (PN). Within each glomerulus, each ORN creates an excitatory connection with one random LN and one random PN. Each PN forms an excitatory connection with one random LN. LNs form connections with all PNs within each glomerulus. Interglomerular communication is mediated by inhibitory connections from each LN to all other LNs and PNs **(Fig. A.3)**.

Each neuron was modelled according to a leaky integrate and fire model (Rozenberg et al., 2019) and evolved according to the membrane potential equation:

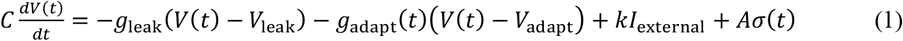

where *C* is the capacitance of the neuron, *g*_leak_ is the conductance of the leaky current, and *g*_adapt_ is that of the adaptive currents. These are modelled according to the equation (Sterratt, 2013):

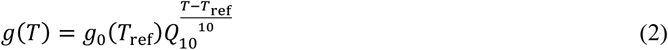

where *T* is the temperature, *V*_leak_ is the reversal potential of the leak current, and *V*_adapt_ is that of the adaptive currents. *I*_external_ accounts for the activity of other neurons or external electrical stimuli. *Aσ*(*t*) is gaussian noise to account for the noisy nature of the neuron scaled by *A*.

Any external signal, coming from an odour or from the external correlated input, was introduced in the model by adding it to the *I*_external_ term of the ORN. The odour receptors were considered a source of noise, adding an external current to the ORNs (Galán, 2006; Joseph J, 2012). In particular, the fraction of odour receptors evolved according to:

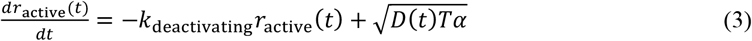

Where α is a normal distribution, *D*(*t*) is computed ad hoc following the *Q*_10_ formalism, *T* is the temperature, and k_deactivating_ is the rate of deactivation of the odour channel.

Once the membrane potential of a neuron reaches *V*_thresh_ a spike is fired and the voltage is decreased to *V*_reset_.

Both the inhibitory and excitatory synapses worked according to the same equation, modifying the conductance of the post-synaptic neuron whenever a spike traversed them:

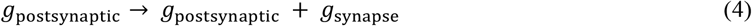

The conductance of the synapses evolved according to an exponential decay equation:

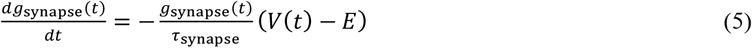

Default-state neuronal and synaptic parameters are summarised in **Tables A.1–A.3**.

The reduction in inhibitory (bot h LN to PN and LN to LN) synapses’ strength has been performed by reducing synapses’ conductance by integer factors, called Synaptic Conductance Reduction (*SCR*) factors. For the network analysis, simulations were run with 10 different coupling strengths, corresponding parameters can be found in **Table A.4**.

The weakly correlated input has been generated to mimic the spontaneous activity, which is also observed in other studies (Galán et al., 2006; Haase, Rigosi, Trona, et al., 2011).

To do so, Poisson processes have been generated as input to individual ORNs, via the Modified Next Reaction Method (MNRM) (Marchetti et al., 2017). For that, spike events are added to the baseline membrane potential:

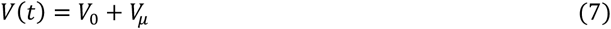

with *V*_*μ*_=1.8 × 10^−2^ mV at random time points:

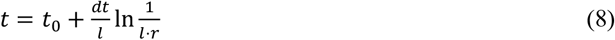

with *l* = 0.5, *r* ~ norm(0,1), *t*_0_ being the previous spike event, and *dt* = 0.2 ms as the simulation time-step.

To control the correlation between ORNs, a template Poisson process was generated by the same method. Correlations between the individual ORN were then introduced by randomly removing events in the individual ORN’s Poisson processes and adding events from the template process with a fixed probability *c =* 0.7. The Poisson processes were then convoluted with an exponential kernel [1/τ exp(*t*/τ)] with *τ* = 2 ms and Kernel width *σ* = 5 *dt* so that they resemble typical electrophysiological traces. The complete computational model was run on an HPC cluster to fully utilize GeNN’s parallelization, simulating 60 s of neuronal activity, and collecting the membrane voltage evolution and spike history for all the neurons.

From the collected data, the same features computed in the experimental settings were extracted. Code and data are available from https://github.com/NeurophysicsTrento/genn-network-model.

## 3. Results

### 3.1. Bee body motion analysis

A frontal camera continuously filmed the motion of the bees’ abdomen while it was head-fixed to the microscopy mount (**Fig. 1a**). A running standard deviation (STD) analysis of the images of the bees’ freely moving abdomen showed epochs of strongly varying signals (**Fig. 1b**) and epochs where bees where nearly motionless (**Fig. 1c**). Averaging the STD over each frame, which produces a time series representing the bee motion over an entire night, indicated two well-separated states of high and low motion activity, with sudden transitions and without extended intermediate levels (**Fig. 1c**). The transitions between the two states were relatively rapid, but not immediate. An analysis of the state transitions (**Fig. 1d**) revealed a characteristic transition time to the sleep state of approximately 50 seconds, while the transition back to the awake state is slightly faster, at around 45 seconds. A real-time thresholding algorithm triggered two-photon brain imaging sessions (**Fig. 1c**, blue vertical lines) whenever such a motion state change was detected. These 2.5 min imaging sessions did not alter the bees’ motion state since the imaging wavelength of 780 nm is outside of the visible spectrum of the bees’ photoreceptors.

### 3.2. Descriptive analysis of neuronal activity time series

The fluorescence signal of a calcium-sensitive dye was recorded in all PNs within an imaging plane passing through the antennal lobe (**Fig. 2a**). A regional homogeneity analysis (**Fig. 2b**) allowed the identification of individual glomeruli and the generation of glomerular masks via image segmentation (**Fig. 2c**). The normalised fluorescence change, averaged over the area of each glomerulus, produced time series (**Fig. 2d,e**) that were considered as a proxy measure for the firing rate of its PNs (Moreaux & Laurent, 2007).

These time series show oscillatory features in numerous glomeruli that represent phases of spontaneous activity. A visual inspection did not immediately reveal clear differences in amplitudes, frequencies, or regularities between recordings from periods of active motion (**Fig. 2d**) and periods of rest (**Fig. 2e**). However, as our subsequent machine learning analysis demonstrates, these differences do exist and can be quantitatively distinguished based on extracted features.

### 3.3. Machine learning analysis

In the machine learning analysis, the target variable to be predicted was whether a sample of neural activity was recorded during an active or rest state. The predicting variables were sets of pre-determined features describing the fluorescence time series. To train a Random forest classifier (RFC), we selected features that focus on three different kinds of properties. First, the overall distribution of neuronal activity, characterised by its moments: the Standard deviation (STD), the Skewness, and the Kurtosis. Second, the complexity of the time series, quantified by its Entropy, the Hurst exponent, and the Detrended fluctuation analysis (DFA). Third, network properties of the glomerular network, quantifying its Betweenness, the Degree, the Efficiency, and the Modularity (see Methods for details).

A qualitative, univariate analysis of the distributions of these features across all subjects and time series (**Fig. 3d**) reveals different outcomes. Some features, such as Kurtosis and Modularity, exhibit highly overlapping distributions in the active and resting states. Others, like the STD and Skewness, display similar distribution shapes but with differing amplitudes. However, the majority of features show noticeable changes in distribution shape: distributions that are monomodal in the active state are bimodal in the sleep state, *e*.*g*. for the Entropy, the Hurst exponents, the DFA. Furthermore, for bimodal distributions, the relative contributions to each mode change considerably, as seen in the network properties Betweenness and Degree. Wilcoxon statistical tests confirmed significant differences between states in all features, except Efficiency and Modularity. Most features showed significant differences also when analysed for each individual bee, although there are exceptions, especially after false-discovery-rate correction (**Table A.5**).

**Figure 3.**
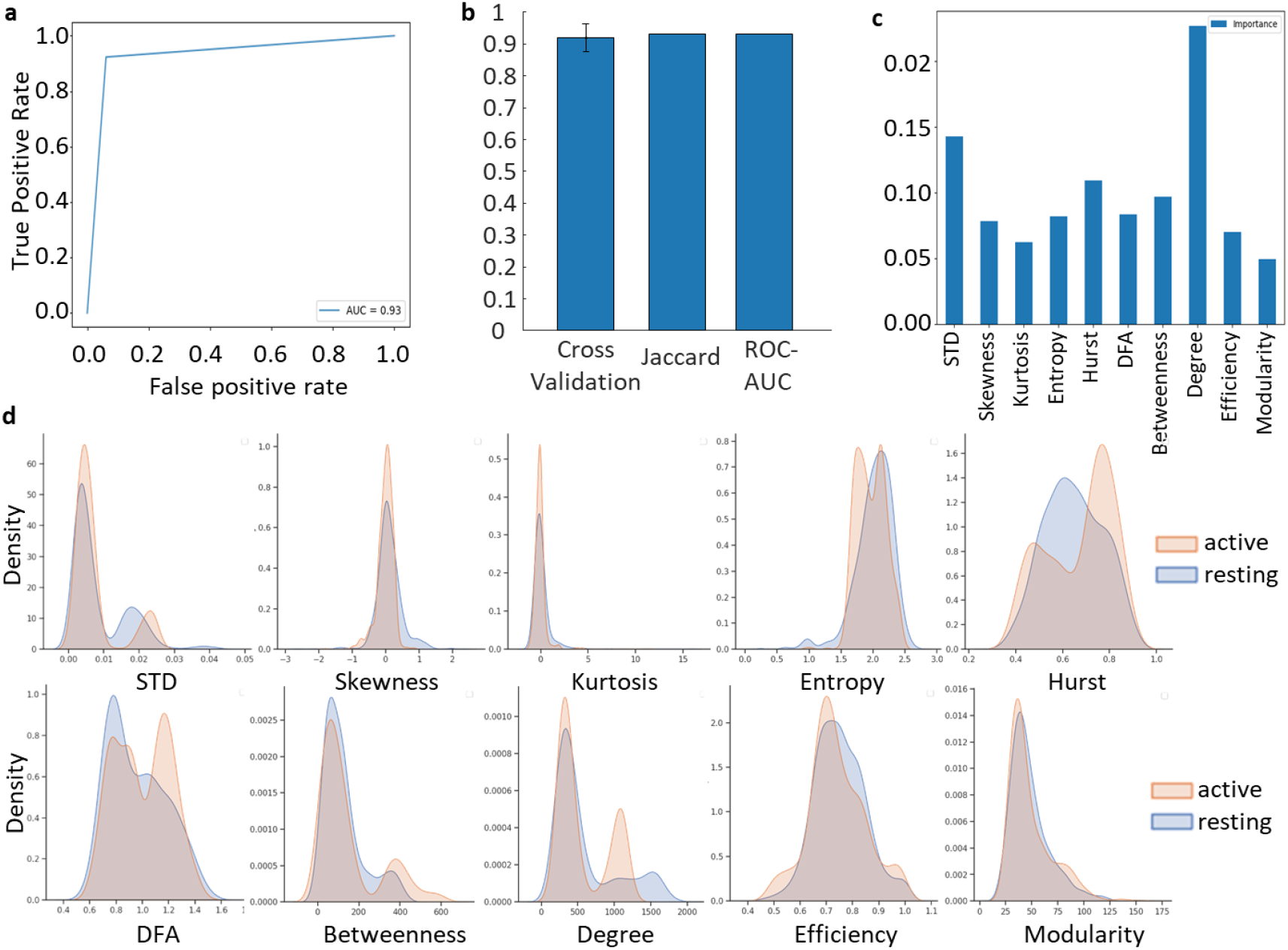
Machine learning classification and feature distributions. (**a**) Receiver-operating characteristic (ROC) curve for the active and rest state classification of the pooled glomerular response data of *n* = 9 subjects for different sensitivity and specificity thresholds; (**b**) Classification results: Cross-validation accuracy, Jaccard index, Area under the ROC curve (ROC-AUC). (**c**) Feature importance analysis comprising moments (Standard deviation (STD), Skewness, Kurtosis), complexity (Entropy, Hurst exponent, Detrended fluctuation analysis (DFA)) and graph theory variables (Betweenness, Degree, Efficiency, Modularity) averaged across glomeruli for all temporal segments and all bees during the active motion phase; (**d**) Distribution of features across all data samples in the active state (orange) and the resting state (blue). The results of the statistical comparisons are given in **Table A.5**.

In addition to analysing the feature distributions within each state, we also examined the distribution of feature differences between states for each glomerulus, for features that are properties of individual glomeruli (**Fig. A.2)**. This allows us to identify whether changes are global, limited to subsets of glomeruli, or if they manifest in opposite directions for different subsets. Notably, the STD shows minimal changes between states across all glomeruli. A case in which most glomeruli exhibit consistent changes in a single direction is the Entropy. For other features, such as Kurtosis, the Hurst exponent, and DFA, the difference distributions are bimodal, with changes occurring in opposite directions across different glomeruli subsets. The RFC results show high distinguishability with an average Cross-validation accuracy of 0.920 ± 0.043; a Jaccard score of 0.932, measuring the intersection between the sets of predicted and true state labels; and an Area under the receiver operating curve (ROC-AUC) of 0.931 (**Fig. 3a,b**).

To exclude the possibility that motion artefacts - although corrected via image registration - are leaking into the brain activity information and could be potentially confounded with sleep vs. awake, another RFC was trained to distinguish between epochs of high vs. low residual brain motion after the registration. Time segments from both body motion states were reordered into two new sets by conducting a median split on the extent of frame displacement detected during a second image registration (**Fig. A.1c**). Parameters and features were chosen identically to the brain activity classification. This classifier failed to distinguish between low and high brain motion states, achieving only near-chance AUC (**Fig. A.1d**). The fact that the motion state could not be distinguished from the time series using this probing procedure suggests the time series contain little information about this variable.

### 3.4. Network analysis

The features analysis suggested substantial differences between global topological features of resting vs. active networks; in particular, the network degree (Fig. 3c) stands out regarding its importance in random forest classification. To further quantify AL network properties, we studied these using standard approaches developed in network science. We computed pairwise correlations among the single glomerular nodes and confronted them with surrogate data sets to quantify differences from spurious correlations. Significant correlations and anti-correlations were observed in most of the glomeruli (Fig. 4). In the active phases, these correlations were relatively small and anti-correlations were present (Fig. 4a), leading to an average Pearson correlation coefficient across all bees of *r* = 0.115 ± 0.032. In the resting epochs, these correlations increased significantly (*t*(16) = −2.9, *p* = 0.0096) and anti-correlations disappeared almost completely (Fig. 4b), leading to an average Pearson’s *r* = 0.265 ± 0.040. These data suggest greater neural synchrony during the sleep state. The influence of residual motion artefacts is negligible, given that the ML classifier was unable to distinguish states based on residual motion (Fig. A.1d).

**Figure 4.**
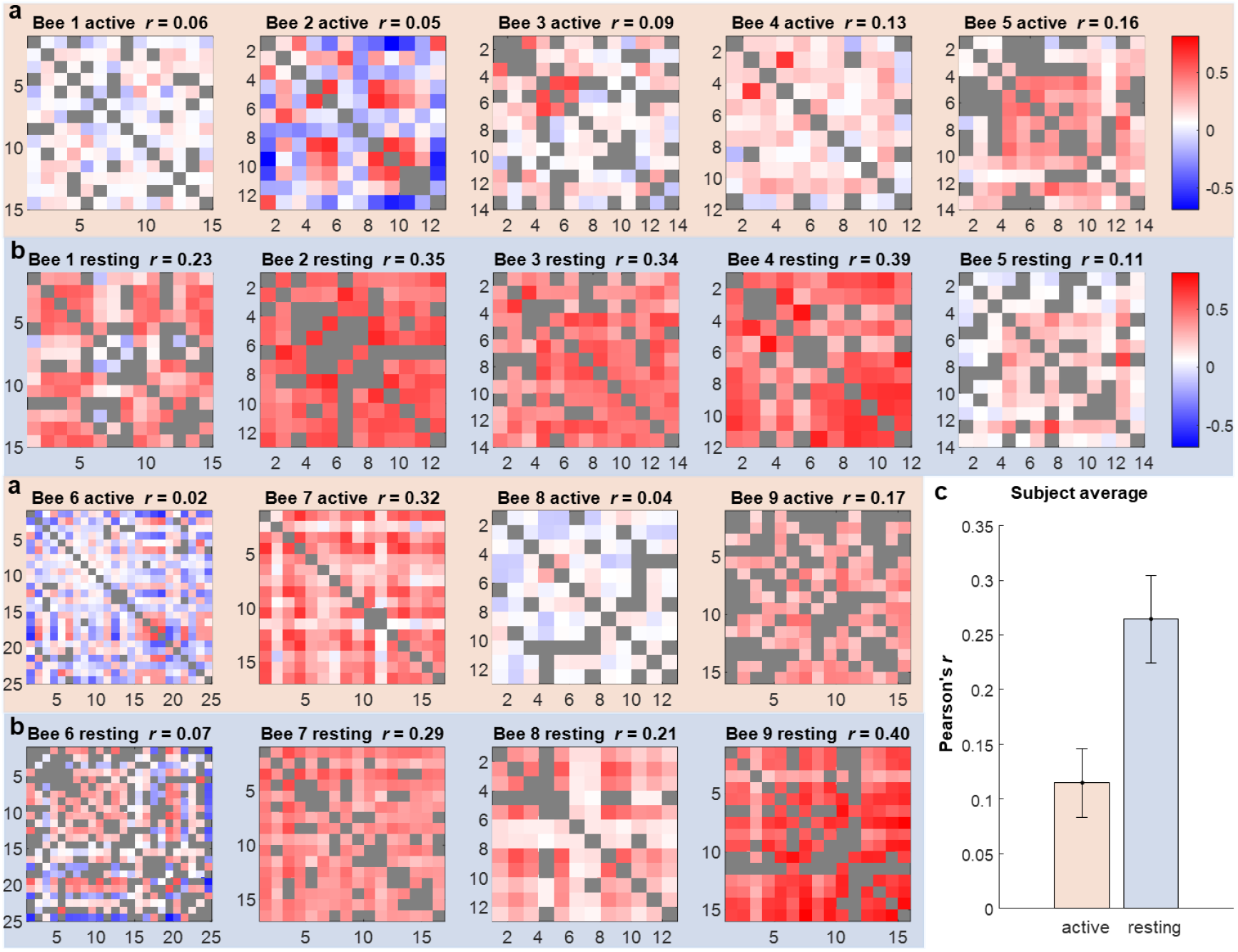
Measured correlations between the activity of individual glomeruli. Correlation matrices for the glomerular activity in *n* = 9 different bees, for the active state (**a**) with an orange background and the resting state (**b**) with a blue background. Pairwise correlations between glomeruli were tested against surrogate data and correlations that did not significantly deviate from chance are marked in grey. For significant correlations, Pearson’s *r* is plotted as a colour code from strong anti-correlations in blue to strong correlations in red. The averaged Pearson’s *r* across the matrix is shown above each matrix. (**c**) Subject-averaged correlation coefficients, with error bars showing the standard deviation. A *t*-test confirms a significant difference in the Pearson correlation between active and resting state (*t*(16) = −2.9, *p* = 0.0096).

### 3.5. Simulations

A computational simulation of the olfactory system was performed to investigate if a spiking neural network (SNN) with variable inhibitory synapse strength could reproduce the observed glomerular coupling patterns. The SNN was modelled as a recurrent spiking neural network following the approach by Scarano *et al*. (Scarano et al., 2023) (**Fig. A.3**). Additionally, a weakly correlated input noise was injected into all receptors to generate weakly time-correlated spontaneous activity, as observed in previous studies (Galán et al., 2006).

The hypothesis tested was that changes in glomerular coupling could be the basis for the experimentally observed change in the correlations between the glomerular outputs, the projection neurons (PNs). We tested this by varying the inhibitory synapse strength in a network of 160 glomerular nodes densely connected by local neurons (LNs) (see methods).

We defined a Synaptic Conductance Reduction (SCR) factor (arbitrary range 1:100) that controls coupling strength between network nodes. Reducing the LN-LN and the LN-PN inhibitory coupling strength by this SCR produced a monotonic increase in glomerular correlations (**Fig. 5a**). Averaging these correlations over all glomerulus pairs shows a sigmoid-shaped growth function of the Pearson’s correlation coefficient (**Fig. 5b**). These correlations saturate at a Pearson’s *r* = 0.37, reaching its half value at *SCR* = 11.7. This suggests that also the experimentally observed effect might be caused by such a reduction in inhibitive coupling. A comparison of the average measured and simulated correlations suggests a synaptic strength of *SCR* ≈ 8 for the active state and *SCR* ≈ 15 for the resting state, which indicates a relative reduction in synaptic coupling strength by a factor of 7.

**Figure 5.**
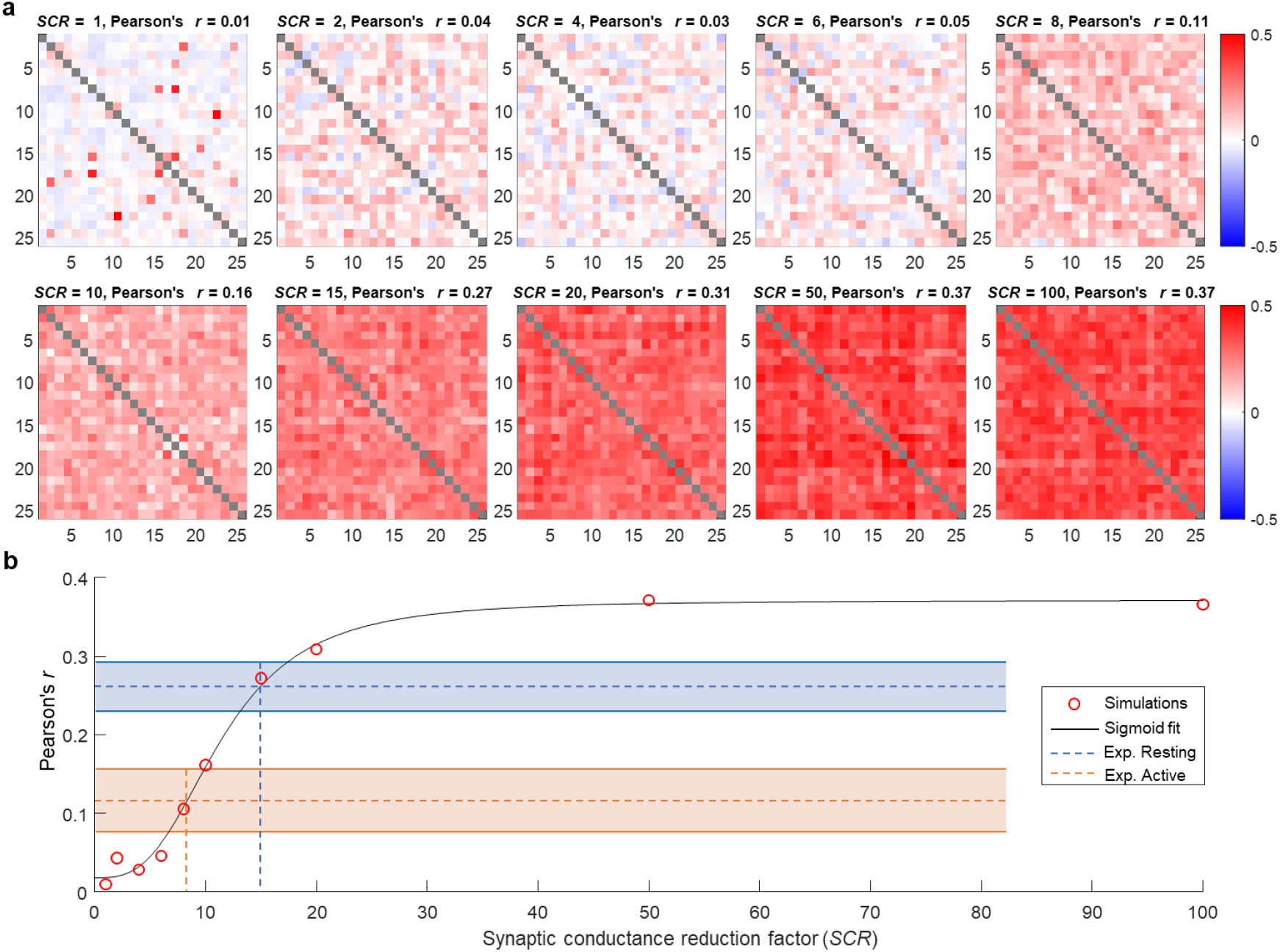
Simulations with varying inhibitory synapse strength: Change in cross-glomerular correlations. **a**) Correlation matrices for the projection neuron (PN) activity in 25 randomly selected glomeruli for simulations with decreasing synaptic conductance (synaptic conductance reduction (*SCR*) factors 1 to 100, corresponding conductances in **Table 1**). (**b**) Dependence of the glomerular correlations on the synaptic conductance: Pearson’s *r* averaged over all PN as a function of the *SCR*. A fitted sigmoid power function gives a saturation level *r* = 0.37 and a half value point *SCR* = 11.5. Dashed horizontal lines and shadowed areas show the experimentally measured correlations and their standard errors, respectively, in the resting state (blue) and in the active state (orange). The vertical lines determine the simulated *SCR* corresponding to the measured correlations in both states.

Next, to test how such changes of inhibitory coupling would influence odour processing in the antennal lobe network, we simulated the input of typical odour signals (Scarano et al., 2023) into the olfactory receptor neurons (ORNs). Odors are simulated by a concentration and set of affinities with a unique Gaussian distribution over all ORNs. This determines the fraction of open channels in each ORN and thus the corresponding change in membrane potential generating the input to each glomerulus. The awake-like network state (*SCR* = 8) produces odour response maps very similar to experimental observations (Paoli et al., 2018), confirming that the simulations closely reproduce the AL functionality. The comparison of the ORN and PN response maps shows the typical characteristics of odour processing in the antennal lobe, namely an increase of contrast and the sparsening of the glomerular code, *i*.*e*. stronger ORN responses are amplified more than weak responses to improve the discriminability of odours (**Fig 6a**). Although the rest-like state PN response maps show that odour processing is still functional, consistent with previous studies (Zwaka et al., 2015), the contrast between highly activated PNs and base level is strongly reduced. The normalised synaptic firing rate falls from 300% in the strongest awake responses (**Fig. 6a**) to 120% in the strongest resting-state responses (**Fig 6b**). A Wilcoxon signed-rank test confirms the highly significant difference between normalised PN firing rates between states (*Z* = −11.0, *p* < 5·10^−28^).

**Figure 6.**
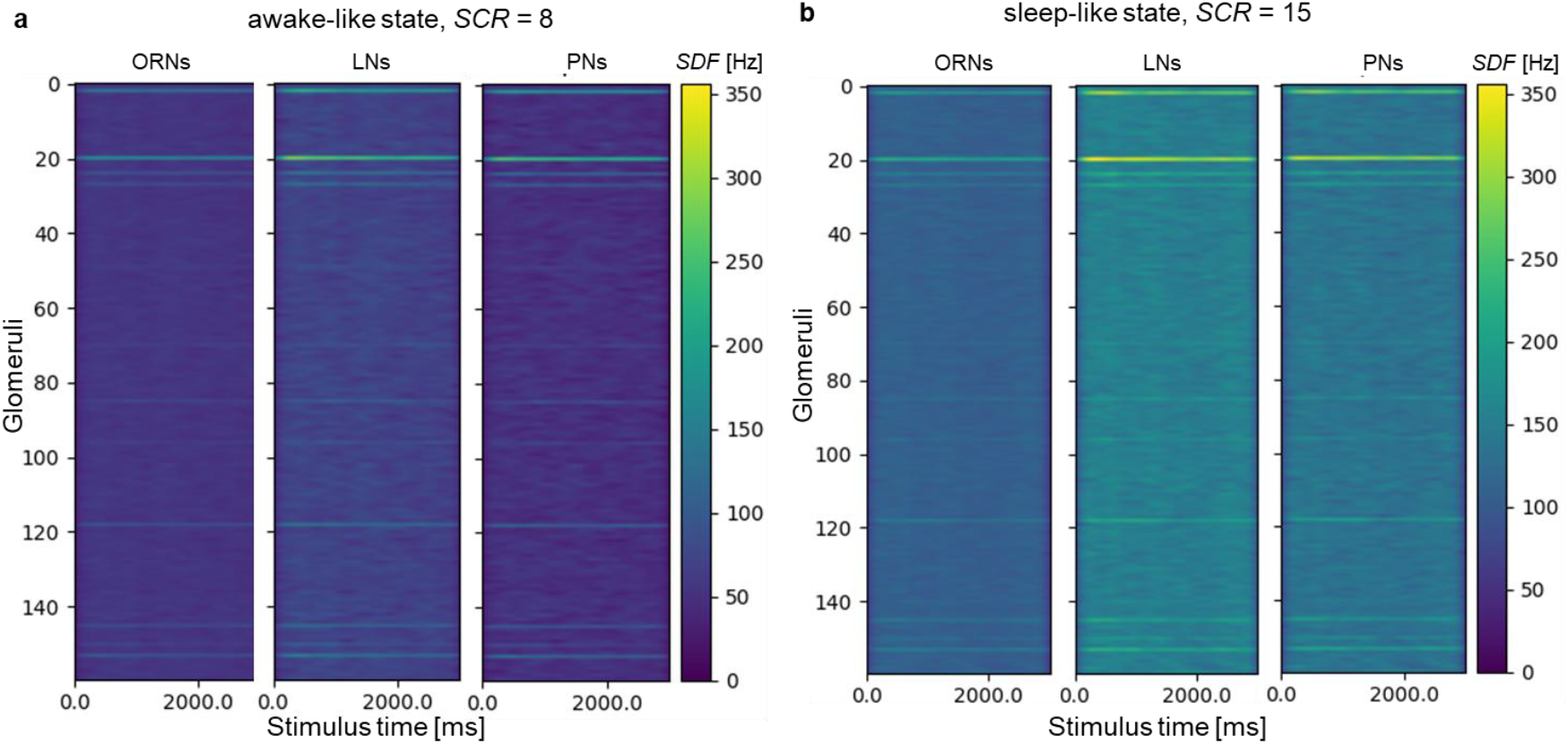
Simulated odour response patterns in the awake- and the sleep-like states. **a**) Spike density function (*SDF*) in the 3 different neuron types: Olfactory receptor neurons (ORNs), Local neurons (LNs), and projection neurons (PNs) for all 160 glomeruli in response to a 3 s odour stimulus simulated in an awake-like state with a synaptic conductance reduction factor *SCR* = 8. **b)** *SDF* response patterns to the same odour stimulus in a sleep-like state with *SCR* = 15. Wilcoxon test of the normalised PN response amplitudes between states confirms a highly significant reduction of contrast (*Z* = −11.0, *p* < 5·10^−28^).

## 4. Discussion

These results represent the first study of long-term activity dynamics in the honey bee brain recorded over entire nights, with analyses of both time-domain and network properties of the measured neuronal activity. Corroborating prior observations, the video recordings of the bees’ body motion show - rather than a continuous variation of motion intensity - switching between two well-distinguished states of low and high motor activity of varying duration from minutes up to hours. The 7 min average duration of the resting states is of the same order as the previously reported 13 min inside the hive (Eban-Rothschild & Bloch, 2008).

Advancing beyond prior behavioural and neural studies, the scope of the current investigation was to determine the network-level neural organisation underlying these states of sleep and wakefulness, in brain systems not directly involved in motor control.

Exploratory data analysis of a sub-network of the antennal lobe shows consistent spontaneous activity across most of its nodes in the form of oscillations with varying frequencies on the order of tenths of Hz. Spontaneous activity with these frequencies has been previously reported, where an increase after odour stimuli was connected to memory formation (Galán et al., 2006).

Among the distribution of selected features describing the active and resting state of the glomerular time series, a common property of the glomerular network stands out: the node degree, i.e. the number of edges connected to each node, shows the strongest deviation between the distributions in the active and in the resting phases.

Examining the differences in features between states at the level of individual glomeruli, we found that there are often subsets of glomeruli in which changes occur in opposite directions. This suggests that the olfactory network is not merely attenuated during sleep, but that alternative processing mechanisms come into play. That said, one feature for which changes almost exclusively go in one direction is the Entropy, which is higher in the rest state in the vast majority of glomeruli. This indicates a higher complexity of the AL output signal during sleep, which again points to sleep-specific neuronal processing. The machine learning analysis proved a clear distinguishability between the multi-glomerular activity states. In fruit flies, a similar approach applied to multichannel electrophysiological recordings was performed focusing on a spectral analysis of the spontaneous activity in the different channels as features, showing clear distinguishability between sleep and awake state patterns and even sleep sub-state discrimination (Jagannathan et al., 2024). The multichannel analysis also covered network properties, which we explicitly operationalised as features.

We found that the network degree strongly contributed to the classification accuracy. The subsequent correlation analysis of the projection neuron response maps showed a highly significant increase in average glomerular cross-correlation and a slight reduction of its variability during the transition into the resting state. Our analyses suggest this is not caused by motion artefacts. Apart from the fact that even the ML classifier was not able to discriminate states based on residual motion, if motion artefacts were a significant factor, they would induce synchronous shifts of all glomeruli, leading to increased correlation during wakefulness.

Similar glomerular cross-correlation levels have been measured in experiments where spontaneous activity was evaluated before and after odour stimulation (Galán et al., 2006). Galán *et al*. found that the change in spontaneous activity synchronisation following odour stimuli matched the reverberations of Hebbian learning theory.

One could hypothesize that the synchronisation changes during sleep observed in our experiments might therefore also be involved in learning and memory consolidation, as these effects have been observed in bees during sleep. One study found that extinction learning was influenced by sleep deprivation, but associative learning was not (Hussaini et al., 2009). Another study also showed the influence of sleep on associative odour learning, as memory could be enhanced by context odour presentation during sleep (Zwaka et al., 2015). Also, navigation memory was found to be consolidated during sleep (Beyaert et al., 2012). The above-mentioned increased complexity in the sleep state could be another indication of memory consolidation processes during sleep.

We then evaluated a possible model of these synchronisation changes by simulating the antennal lobe at a realistic size via a recurrent spiking neural network (Scarano et al., 2023). In order to reproduce the experimentally observed spontaneous activity, weakly correlated Gaussian noise was fed into the olfactory neurons. The role of such a noise input in the reproduction of important neuronal mechanisms was previously shown in a simulation of learning and memory formation in insects via a neural network model which, besides antennal lobes, included mushroom bodies and lateral horns (Arena et al., 2012). Also there, Gaussian noise, which was used to simulate ORN input during sleep, gave rise to pattern formation in the MBs, as would be expected during overnight memory consolidation.

In our antennal lobe model, the transition between awake and sleep-like states required only a single parameter to be changed: the inhibitory synaptic conductance that couples LNs to LNs and LNs to PNs. A decrease on the order of 10 caused a change in connectivity as observed experimentally. This is consistent with the human sleep theory of synaptic homeostasis, which suggests that synaptic strength is downscaled to a baseline level that is energetically favourable and beneficial for learning and memory (Tononi & Massimini, 2008).

Our simulations also revealed the differences in odour processing between sleep and awake network states. In the awake state, the model demonstrates a clear increase of contrast in the activation patterns but also the characteristic differential gain control, which more strongly amplifies highly activated ORNs compared to weakly activated ones (Root et al., 2008). This leads to a sparsening of the odour code, a mechanism that improves odour distinguishability. This reduction in odour processing might then indeed lead to a reduced specific sensitivity to certain odours, which however should be overcome by stronger stimulation; both are characteristics that define sleep (Helfrich-Förster, 2018). The general sensitivity to specific odours during sleep, which our model confirms, was shown by the above-mentioned memory consolidation experiments (Zwaka et al., 2015).

In *Drosophila*, it was previously reported that rest phases demonstrate sleep-like characteristics, including reduced responsiveness to sensory stimuli (Hendricks et al., 2000). Calcium imaging experiments in the mushroom bodies (MBs) showed reduced baseline activity during sleep phases (Bushey et al., 2015). Our results from the antennal lobes, which are upstream of the mushroom bodies, do not show a general reduction of the activity but rather a change in information integration: the input signal is forwarded without the contrast enhancement that is observed in the awake state. This suggests that the observed reduction in the response patterns in the higher brain centers is produced by a decreased integration of the peripheral responses, rather than reduced responsiveness of receptors or primary neurons.

Our simulations suggest that reduced inhibitory coupling causes the observed network property changes, possibly via changes in the neurotransmitter concentration. For the honey bee, this neurotransmitter is likely GABA, which is dominant in the antennal lobe (Schäfer & Bicker, 1986). In contrast to *Drosophila* (Huang et al., 2010), excitatory LNs have not been found in bees.

The involvement of GABA modulation in sleep has been observed in numerous studies in a variety of species from *Drosophila* (Agosto et al., 2008) to humans (Gottesmann, 2002). In *Drosophila* a reduction of GABA release reduces total sleep (Agosto et al. 2008) as does a knock-down of GABA receptors in *Drosophila* circadian pacemaker neurons (Chung et al. 2009).

In humans, however, the opposite effect has also been reported: while an increased cortical GABA concentration was found initially during Non-rapid eye movement (NREM) sleep, it decreased progressively after 1 h of recovery sleep and was significantly lower during Rapid eye movement (REM) sleep (Vanini et al., 2012). A distinction between different sleep types could not be made based on our limited behavioural data. However, the bimodal distributions that we observed in several features of our time series suggest that further sub-states might be present. Also in Drosophila, a distinction between wake-like sleep phases and less active deep sleep stages was made by calcium imaging of neuronal activity (Tainton-Heap et al., 2021).

This supports the idea that the neuroregulatory mechanisms responsible for sleep-like states are, at least to some degree, conserved throughout evolution (Parisky et al. 2008).

The reduced information processing in the sleep state shown in our simulations is consistent with a reduced transmission ratio between the peripheral sensory organs and the central brain observed in humans (Coenen & Drinkenburg, 2002).

The alignment of our findings with previous studies suggests that this new method of directly observing network activity modulation during sleep at the single-neuron level offers valuable potential for future research. This approach may provide more detailed mechanistic insights into the underlying processes. Beyond monitoring single-glomerular activity, exploring the role of neuronal connectivity within networks could be particularly useful for understanding how sleep architecture is formed. This includes identifying sleep states at the network level and understanding the transitions between them, as well as how circadian and homeostatic sleep regulation mechanisms affect network states. Achieving this will require more detailed observations of sleep behaviour in bees. So far, we have only assessed whole-body motion, but focusing on antennal movement could provide further insights into potential sleep sub-states, as suggested by previous studies on body and antenna posture in honeybees (S Sauer et al., 2003) and posture and neuronal response patterns in *Drosophila* (Jagannathan et al., 2024).

Combining learning experiments with imaging of sleep-dependent neuronal alterations could deepen our understanding of the connection between sleep and long-term memory formation. While this relationship is well-established by behavioural studies in humans (Walker, 2009) and other species (Vorster & Born, 2015), the neural mechanisms are largely unknown. Comparing findings from this animal model with human sleep studies could offer new evolutionary insights into the function and significance of sleep (Keene & Duboue, 2018; Rößler & Klein, 2024). This research may also provide translational results, potentially contributing to new approaches for diagnosing and treating sleep disorders (Toth & Bhargava, 2013).

## 5. Conclusion

To summarise, this is the first long-term brain imaging study in honey bees during sleep and wakefulness. It reveals different neuronal dynamics in the antennal lobes at the network level. Active and resting states are clearly distinguishable, both in terms of body movement and neuronal activity, with resting states showing increased network synchrony and complexity. Simulations suggest that reduced inhibitory coupling, likely due to GABAergic modulation, underlies these changes. These findings provide novel insights into sleep-related changes in network properties of the invertebrate brain, revealing the neural basis for reduced sensory information processing during rest. Our findings suggest a conserved role of inhibitory modulation in sleep regulation, drawing parallels to synaptic homeostasis models in vertebrates. These results highlight the utility of honey bees as a model system for studying fundamental sleep mechanisms at the neuronal network level and open the door for understanding sleep-related processes, such as their relationship to memory formation.

## Acknowledgements

We acknowledge financial support from the University of Trento Strategic Project Brain “Network Dynamics (BRANDY)”.

## Appendix

**Figure A.1. Motion correction**

The between-frame motion before and after the motion correction via an HMM algorithm. Above: Displacements in each time segment along the *x*- and *y*-axis for three example bees (**a**) before, (**b**) after motion correction. **(c)** Machine-learning analysis for the residual brain motion classification: ROC curves. **(d)** Feature importance analysis for the classification.

**Figure A.2. Feature difference distribution**

Features used for machine learning classification analysed for individual glomeruli: normalised count of glomeruli for the occurring difference values: awake - resting state. Shown are mean +/- std after averaging histograms of individual bees.

**Figure A.3. AL model structure**

Structure of the antennal lobe in the computational simulation framework, shown for 3 glomeruli. Excitatory connections are made from ORNs to PNs and LNs, and from PNs to LNs of the same glomerulus. Inhibitory dense connections are found from LNs to PNs and LNs between different glomeruli [from (Scarano et al., 2023)].

**Figure A.4. Image post-processing steps**

**(a)** Example of raw data in a single time window, **(b)** after application of a smoothing kernel, **(c)** after mean normalisation, **(d)** after thresholding.

**Equations A.1-A.10: Feature formulas**

- **Betweenness:** The Node betweenness centrality *C*_B_ is the fraction of all shortest paths in the network that contain a given node *v*. Nodes with high values of betweenness centrality participate in a large number of paths.

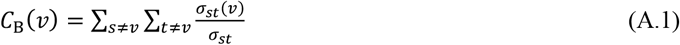

where *σ*_*st*_ is the number of shortest paths from vertex *s* to vertex *t*, and *σ*_*st*_(*v*) is the number of shortest paths from *s* to *t* that go through vertex *v*. The feature is then given by the averaged *C*_B_ across nodes.
- **Degree:** The Node degree *C*_D_ is the number of edges connected to the node *v*.

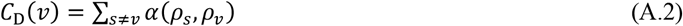

where *α* is a scaling factor multiplied by the number of edges adjacent to node *v*.
- **Efficiency** *C*_E_ is the average of inverse shortest path length, and is inversely related to the characteristic path length.

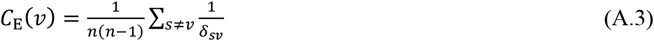

where δ is the characteristic path length.
- **Modularity** *C*_M_ is a statistic that quantifies the degree to which the network may be subdivided into such clearly delineated groups.

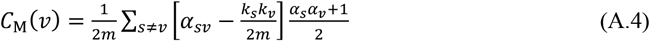

where 2*m* is the number of half edges, α is the expected number of edges, and *k* is the number of nodes.
- **Entropy:** The sample entropy *E* measures the complexity of a time series, based on approximate entropy.

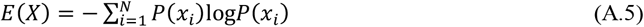

where *P* is the discrete probability of each time point.
- **Hurst exponent** *H* is a measure of the long-term memory of a time series. It can be used to determine whether the time series is more, less, or equally likely to increase if it has increased in previous steps.

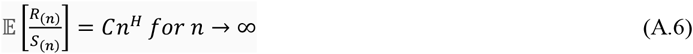

where 𝔼 is the expected value, *n* is the time span of the observation, *S* is the series sum, *R* is the range, and *C* is a constant.
- **Detrended fluctuation analysis (DFA)** *F* is similar to the Hurst exponent but can be applied to non-stationary processes (whose mean and/or variance change over time).

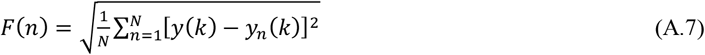

where *N* is the length of the time series, and y_n_ is the piecewise sequence of straight-line fits of the local trends using least squares regression.
- **Standard deviation (STD)** *σ* is the second moment of a distribution, measuring the variation or dispersion of a set of time-series values.

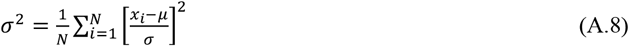

where *µ* is the mean of the distribution and *N* is the total number of samples.
- **Skewness** *μ*_3_ is the third moment, measuring its asymmetry of the distribution of time-series values.

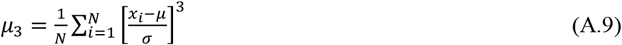

where *µ* is the mean of the distribution, *N* the total number of samples, and *σ* the standard deviation.
- **Kurtosis** *μ*_4_ is the fourth moment, measuring how much the tails of a distribution differ from the tails of a normal distribution of time-series values.

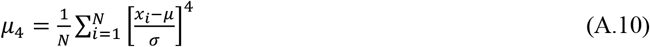

where *µ* is the mean of the distribution, *N* the total number of samples, and *σ* the standard deviation.

**Table A.1:** Parameters for the neurons. (ORN, LN, PN have identical properties, except for the ORN noise input)

| Parameter | Value | Parameter | Value | Parameter | Value |
| --- | --- | --- | --- | --- | --- |
| $C$ | 1 mF | $g_{\text{leak}0}$ | 0.01 S | $k_{\text{deactivating}}$ | 0.0025 |
| $V_{\text{reset}}$ | −70 mV | $g_{\text{adapt}0}$ | 0.00015 S | $T_{\text{ref}}$ | 36°C |
| $V_{\text{thresh}}$ | −40 mV | $r_{\text{scale}}$ | 10 | $T$ | 30°C |
| $V_{\text{leak}}$ | −60 mV | $\tau_{\text{adapt}}$ | 1000 ms | $Q_{10}$ | 1.1 |
| $V_{\text{adapt}}$ | −70 mV | $D$ | $3 \cdot 10^{-5}$ | $A\sigma$ | $1.4/\sqrt{dt}$ |

**Table A.2:** Parameters for the excitatory synapses.

| ORN to PN |  | ORN to LN |  | PN to LN |  |
| --- | --- | --- | --- | --- | --- |
| Parameter | Value | Parameter | Value | Parameter | Value |
| $g$ | 0.008 S | $g$ | 0.008 S | $g$ | 0.001 S |
| $E$ | 0 mV | $E$ | 0 mV | $E$ | 0 mV |
| $\tau$ | 10 ms | $\tau$ | 10 ms | $\tau$ | 10 ms |

**Table A.3:** Parameters for the inhibitory synapses.

| LN to PN |  | LN to LN |  |
| --- | --- | --- | --- |
| Parameter | Value | Parameter | Value |
| $g$ | $5.5 \cdot 10^{-5}$ S | $g$ | $2.0 \cdot 10^{-5}$ S |
| $E$ | −80 mV | $E$ | −80 mV |
| $\tau$ | 20 ms | $\tau$ | 20 ms |

**Table A.4:**
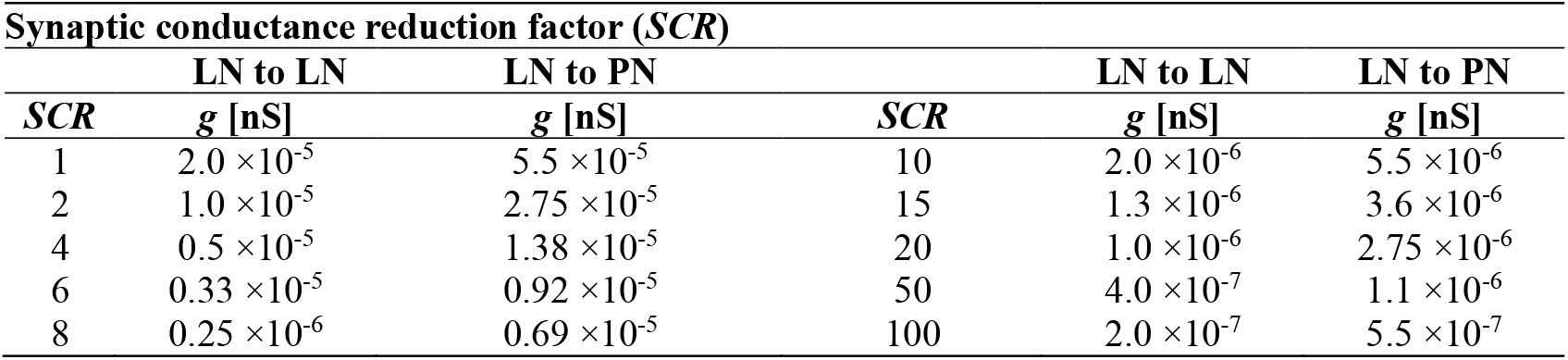
Absolute synaptic conductances corresponding to the synaptic conductance reduction factors.

**Table A.5:**
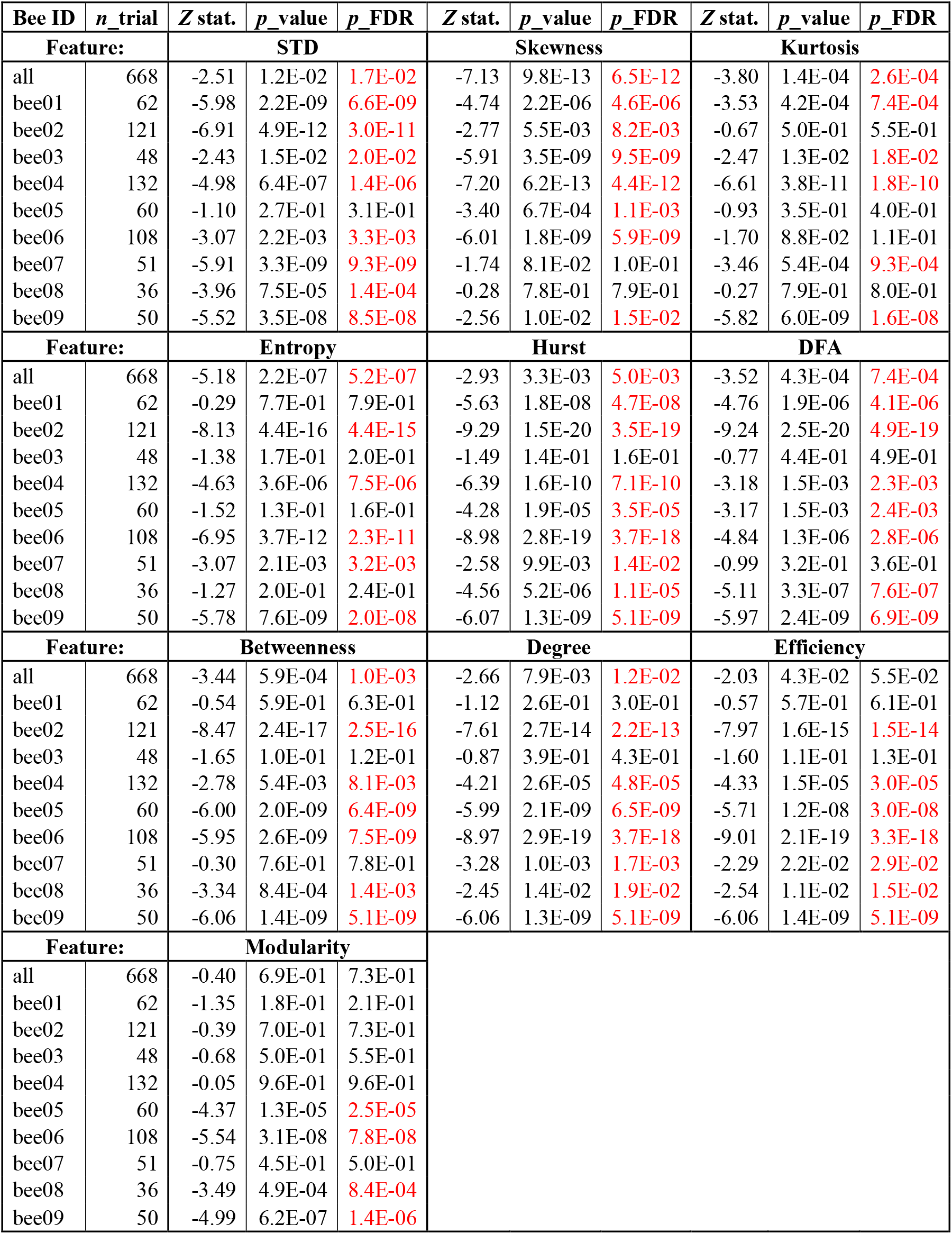
Feature distribution differences between sleep and awake.

