## Supplementary figures and images for "Neuronal correlates of sleep in honey bees"

### Supplementary Fig. A1

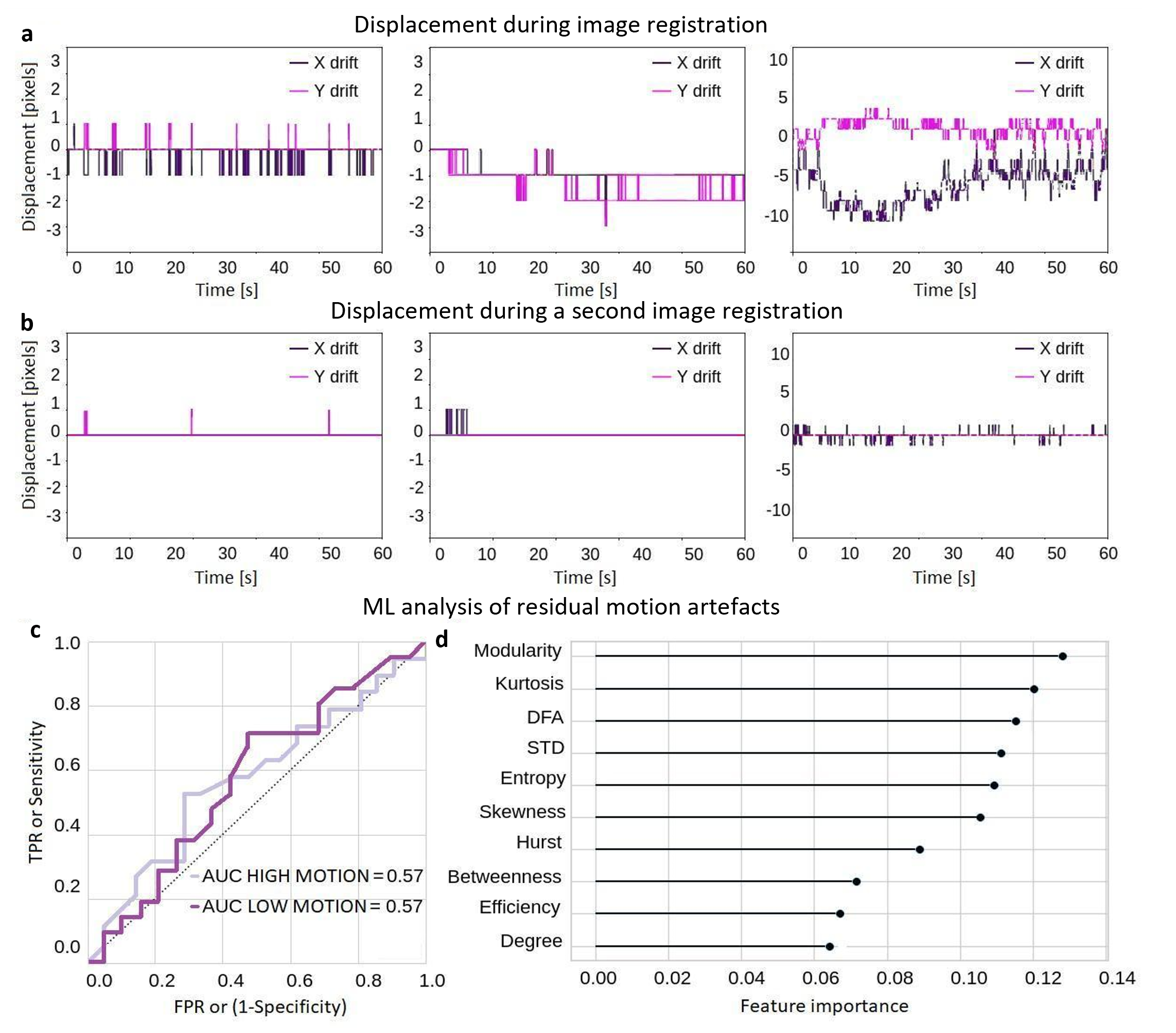

### Supplementary Fig. A2

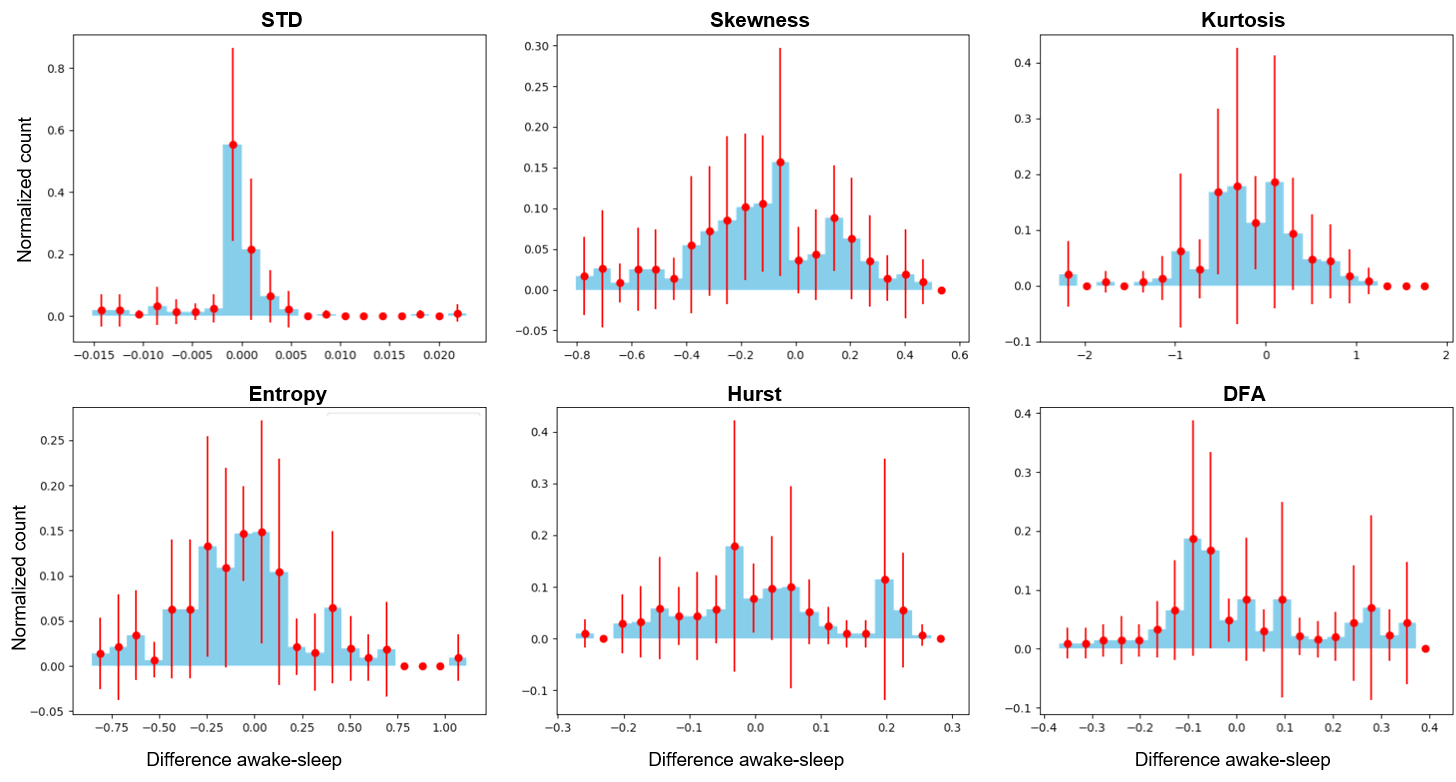

### Supplementary Fig. A3

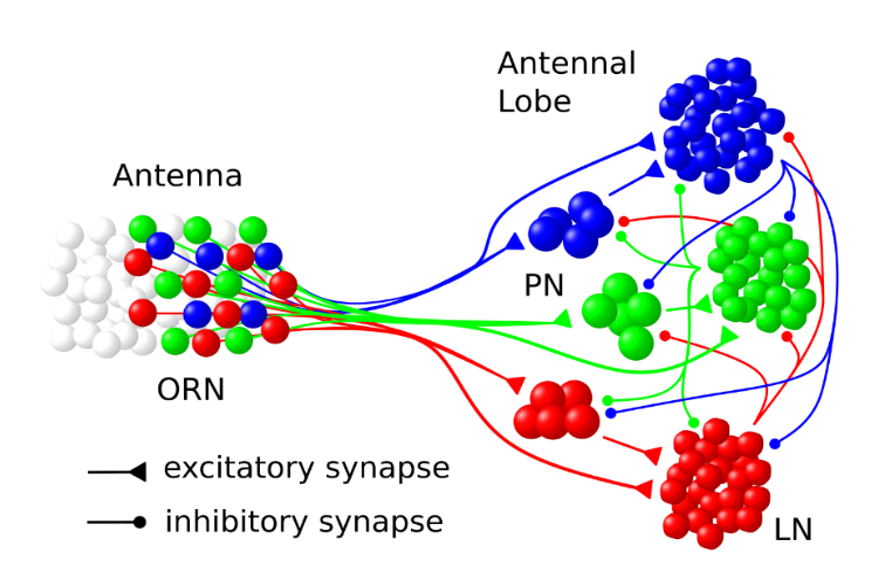

### Supplementary Fig. A4

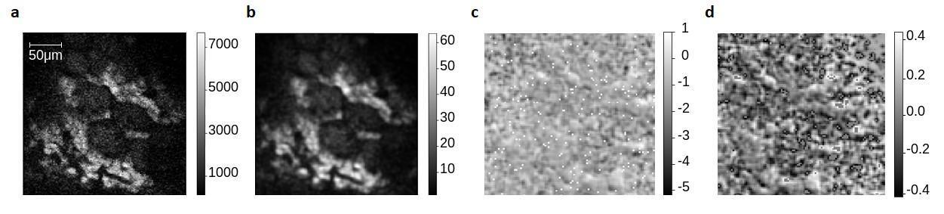
